# Bacterial flagellar motor PL-ring disassembly Sub-complexes are widespread and ancient

**DOI:** 10.1101/786715

**Authors:** Mohammed Kaplan, Michael J. Sweredoski, João P.G.L.M. Rodrigues, Elitza I. Tocheva, Yi-Wei Chang, Davi R. Ortega, Morgan Beeby, Grant J. Jensen

## Abstract

The bacterial flagellar motor is an amazing nanomachine. Understanding how such complex structures arose is crucial to our understanding of cellular evolution. We and others recently reported that in several Gammaproteobacterial species, a relic sub-complex comprising the decorated P- and L-rings persists in the outer membrane after flagellum disassembly. Imaging nine additional species with cryo-electron tomography, here we show that this sub-complex persists after flagellum disassembly in other phyla as well. Bioinformatic analyses fail to show evidence of any recent horizontal transfers of the P- and L-ring genes, suggesting that this sub-complex and its persistence is an ancient and conserved feature of the flagellar motor. We hypothesize that one function of the P- and L-rings is to seal the outer membrane after motor disassembly.

## Introduction

The bacterial flagellar motor is one of the most famous macromolecular machines, made up of thousands of protein subunits that self-assemble in a highly-synchronized manner into a motor, a flexible hook, and a long extracellular filament that rotates in a propeller-like fashion to move the cell (1). The process of how these different parts assemble has been studied extensively using different biophysical and biochemical methods (2–7). These studies have resulted in the current “inside-out” model which starts with the assembly of an inner-membrane-embedded type III secretion system (T3SS) export apparatus, a membrane/supramembrane (MS) ring, a cytoplasmic switch complex (aka C-ring) and a periplasmic rod which connects the MS ring to the extracellular hook. The P- (peptidoglycan) and L- (lipopolysaccharide) rings surround the rod in the periplasm and are thought to act as a bushing during rotation. Finally, the hook is connected by junction proteins to the long filament. While almost all species have this conserved core, different species can have additional cytoplasmic, periplasmic and extracellular components (8–12). For example, in some species (like *Vibrio* spp.) the P- and L-rings are decorated by five proteins (MotX, MotY, FlgO, FlgP and FlgT) (13, 14). In other species, like *Legionella pneumophila* and *Pseudomonas aeruginosa*, the P-ring is decorated by a ring formed by MotY (9).

Much less is known about the process of flagellar disassembly, though it is known that *Caulobacter crescentus* ejects its flagellum and pili at a specific stage of its life cycle (15). We and others also recently reported that different Gammaproteobacteria species lose their flagella when starving or due to mechanical stress (7, 16–18). Interestingly, in situ imaging using cryo-electron tomography (cryo-ET) showed that this disassembly process leaves an outer-membrane associated relic sub-complex consisting of the decorated flagellar P-(peptidoglycan) and L-(lipopolysaccharide) rings (referred to henceforth as PL sub-complexes). These PL sub-complexes plug the hole in the outer membrane that might otherwise be present after the flagellum disassembles. However, it remains unclear whether these PL sub-complexes only persist in Gammaproteobacteria or if the phenomenon is more widespread.

Here, using a combination of cryo-ET (19) and subtomogram averaging (20, 21) we show that the PL sub-complex persists in nine additional bacterial species including *Vibrio cholerae, Vibrio harveyi* and *Vibrio fischeri* (sheathed Gammaproteobacteria); *Hyphomonas neptunium*, *Agrobacterium tumefaciens, Caulobacter crescentus* (Alphaproteobacteria); *Hylemonella gracilis* (Betaproteobacterium); *Campylobacter jejuni* (Epsilonproteobacterium); and *Acetonema longum* (Firmicutes). Bioinformatics analyses further show that the P- and L-ring genes are ancient and diverged separately in each species (were not recently transferred horizontally). Together these results suggest that the outer-membrane-sealing role of the PL sub-complexes is ancient and widely conserved.

## Results

To examine the generality of PL sub-complex persistence, and how the presence of a membranous sheath surrounding the flagellum might affect this process, we used cryo-ET to image nine additional bacterial species from four new classes (Fig. 1). All previously described PL sub-complex subtomogram averages have been of species with unsheathed flagella: *Shewanella oneidensis*, *Legionella pneumophila*, *Pseudomonas aeruginosa, Salmonella enterica* and *Plesiomonas shigelloides* (7, 16, 17) (Fig. S1). All of these feature a crater-like structure in the outer membrane (see examples in Fig. S1), sealed across the bottom by either the P- or L-ring proteins or additional, as-yet-unidentified molecules. This presumably is to avoid an ~ 20 nm pore in the outer membrane, which might be detrimental to the cell. For this reason, we were first interested in whether there would be similar discontinuities in the outer membrane in species with sheathed flagella (in which the flagellum does not always penetrate the outer membrane). Images of individual PL sub-complexes in *V. cholerae* and *V. fischeri* have been published (16), but no subtomogram averages are available. Thus we first imaged the three Gammaproteobacterial species *V. cholerae, V. harveyi* and *V. fischeri*, whose flagella are sheathed. As expected, we observed that the outer membrane of all three *Vibrio* species bent and extended to sheath the micrometers-long extracellular flagellar filaments (Fig. 2 a-c). At the base of these filaments, flagellar motors were clearly visible. Next to the fully-assembled motors, we occasionally observed PL sub-complexes (Fig. 2 d-f). Sub-tomogram averages of these sub-complexes confirmed that they indeed consist of the embellished P- and L-rings (Fig. 2 g-i). In contrast to the structures previously observed from unsheathed flagella, the *Vibrio* spp. structures reported here exhibit an intact, convex outer membrane layer across the top (Fig. 2 g-i). The bottom of the PL sub-complex is still plugged, however (Fig. 2g-i, yellow arrows), raising the question of why.

**Figure 1:**
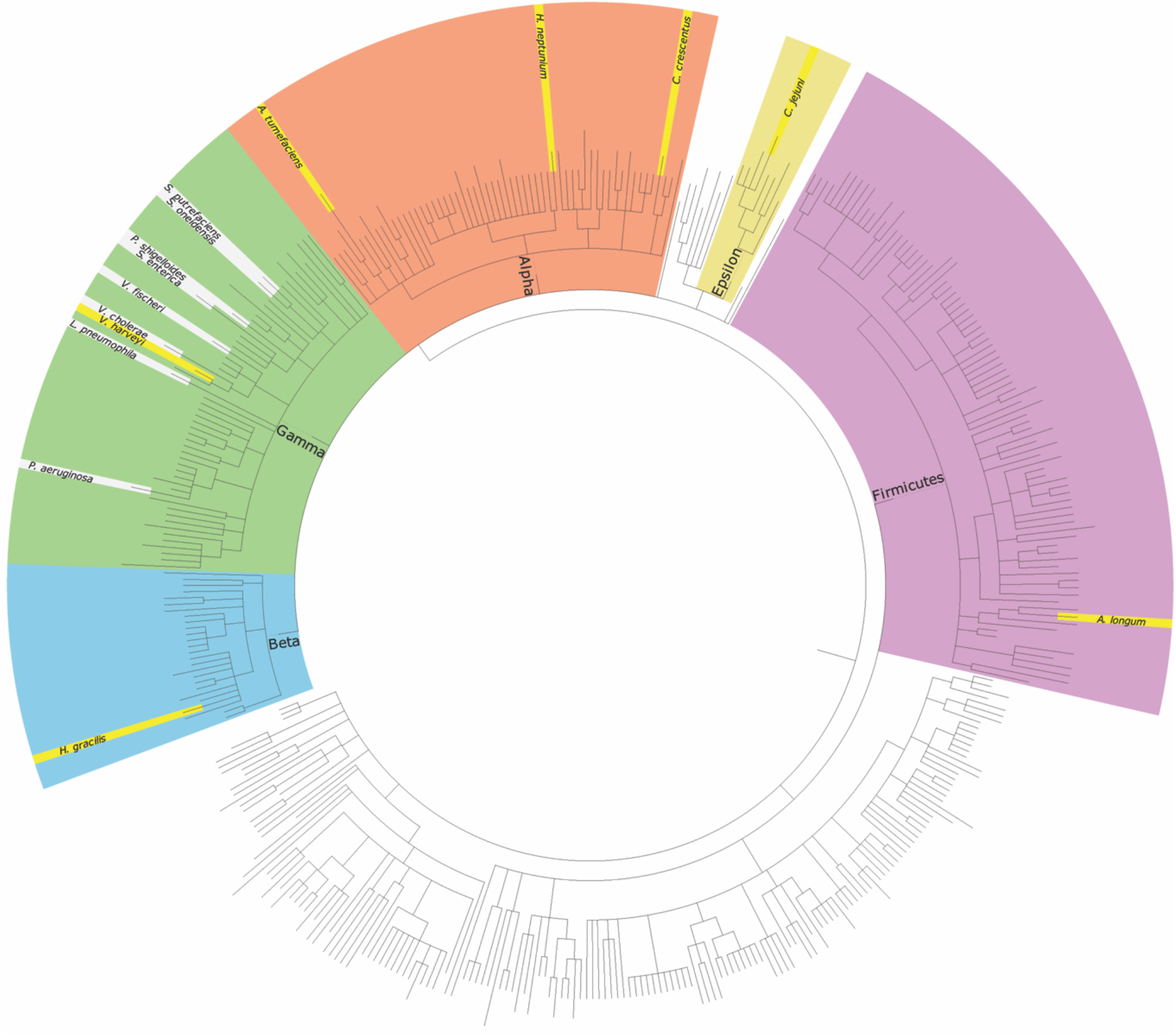
A taxonomic tree of representative bacterial species. The species where PL sub-complexes were previously reported are highlighted in grey (all in the Gammaproteobacteria class) while species with PL sub-complexes identified in this study are highlighted in yellow.

**Figure 2:**
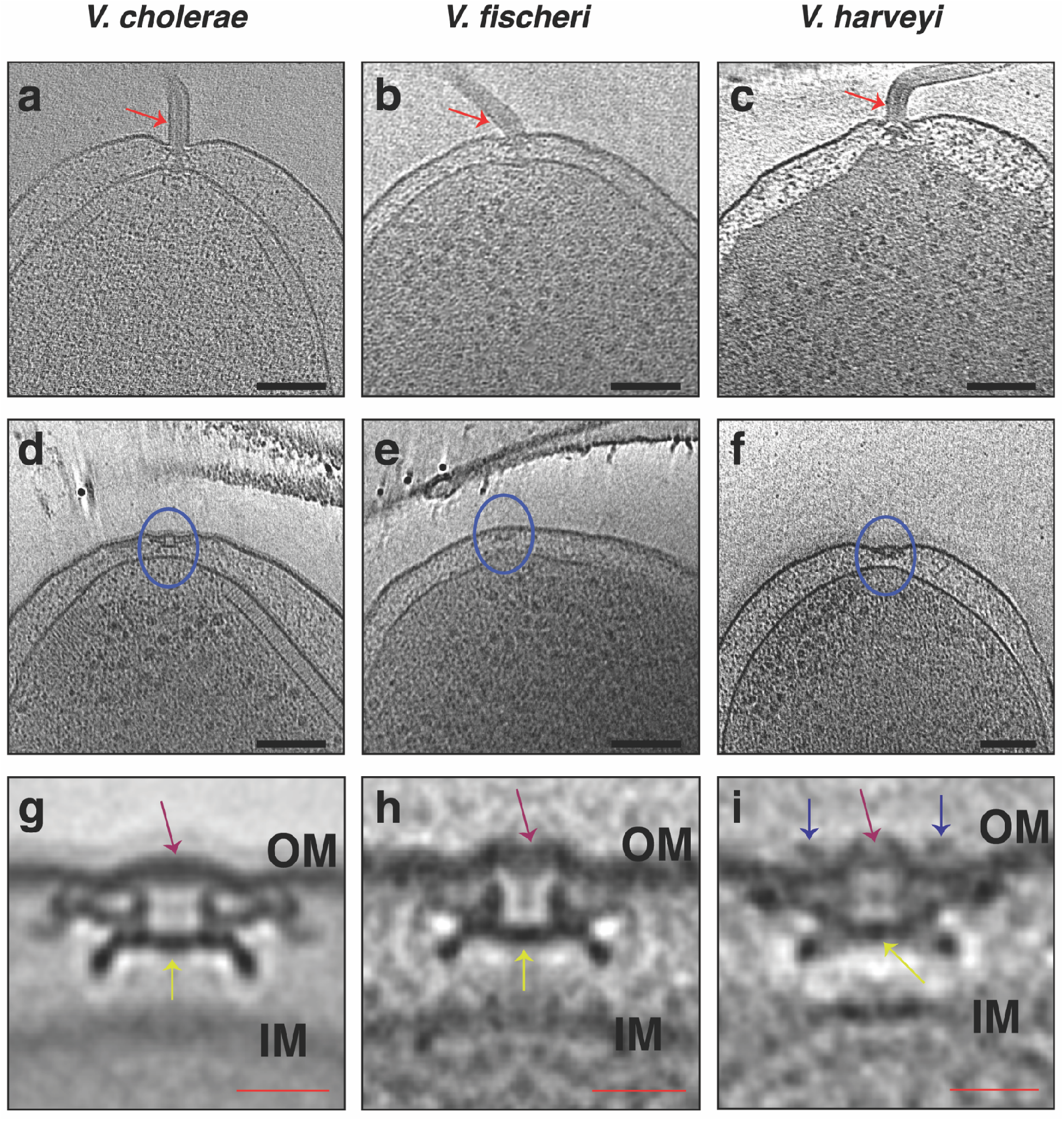
Cryo-ET of the sheathed Gammaproteobacteria *Vibrio* species. **a, b and c**) Slices through electron cryo-tomograms of *V. cholerae*, *V. fischeri* and *V. harveyi*, respectively, highlighting the presence of a single polar sheathed flagellum in the three species (red arrows). Scale bars are 100 nm. **d, e and f**) Slices through electron cryo-tomograms of *V. cholerae, V. fischeri* and *V. harveyi*, respectively, highlighting the presence of flagellar disassembly PL sub-complexes (blue circles). Scale bars are 100 nm. **g, h and i**) Central slices through sub-tomogram averages of PL sub-complexes in *V. cholerae*, *V. fischeri* and *V. harveyi*, respectively. Purple arrows highlight the presence of intact outer membrane (OM) above the PL sub-complexes. Yellow arrows indicate the proteinaceous plug inside the P-ring. Blue arrows in (i) highlight the presence of an extracellular ring density in the average of *V. harveyi*. Scale bars are 20 nm.

In addition, the structure of the PL sub-complex in *V. harveyi* has an extracellular ring located just above the outer membrane (Fig. 2 I, blue arrows). Such a ring is also present in the fully-assembled sheathed motor also (Fig. S2, blue arrows). However, while the diameter of this ring is 30 nm in the PL sub-complex, it has a diameter of 36 nm in the fully-assembled motor suggesting that this ring collapses upon flagellar disassembly. The presence of extracellular rings has previously been described in the unsheathed motor of *S. oneidensis* (9), and the sheathed motor of *Vibrio alginolyticus* (22). Importantly, the structure of the PL sub-complex from *S. oneidensis* has an extra density located just at the membranous discontinuity resulting from disassembling the flagellum (Fig. S1 a). This density in *S. oneidensis* may also be due to the collapse of the extracellular ring present in the full motor.

After this comparison of the PL sub-complexes in the sheathed and unsheathed flagella of Gammaproteobacteria, we were interested in whether PL sub-complexes are specific to Gammaproteobacteria or present in other classes in the Proteobacteria phylum. We therefore examined five more species: *Hyphomonas neptunium*, *Agrobacterium tumefaciens*, and *Caulobacter crescentus* (Alphaproteobacteria, (Fig. 3 a-t)); *Hylemonella gracilis* (Betaproteobacterium, (Fig. 4 a-d)); and *Campylobacter jejuni* (Epsilonproteobacterium, (Fig. 4 e-f)). PL sub-complexes were observed in all of these species with the characteristic bend in the outer membrane and a plugged base similar to their Gammaproteobacterial counterparts. In *C. jejuni*, an inner-membrane-associated sub-complex of the flagellar motor (constituting the MS- and C-rings, the export apparatus and the proximal rod) was present in the vicinity of the PL-sub-complex in a pattern reminiscent to what has recently been reported in *L. pneumophila* (7) (see movie S1 and Fig. S3).

**Figure 3:**
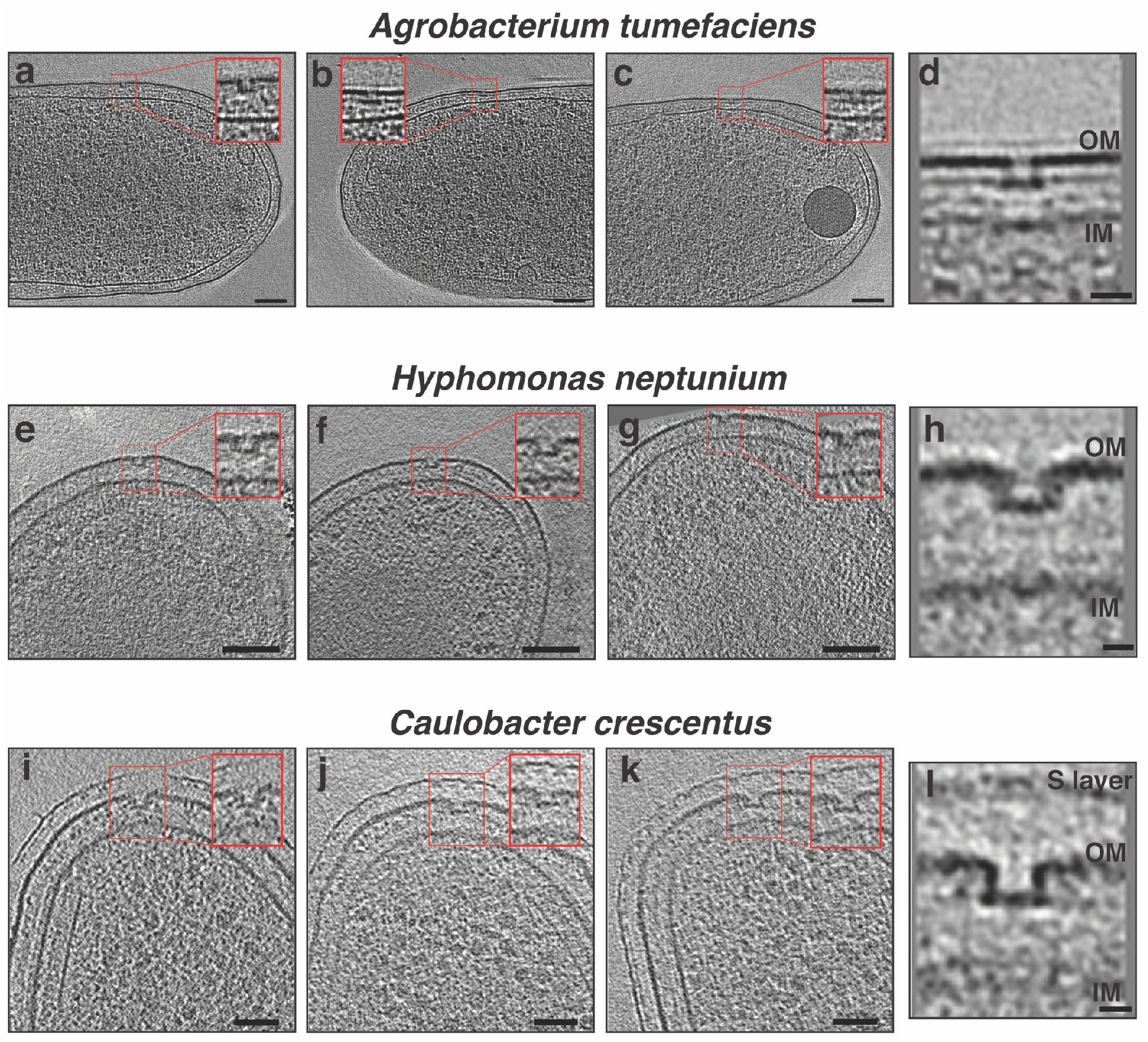
Cryo-ET of the Alphaproteobacteria species. **a, b and c**) Slices through electron cryotomograms of *A. tumefaciens* highlighting the presence of flagellar disassembly PL sub-complexes with zoom-ins of these sub-complexes present in the red squares. Scale bars are 100 nm. **d**) Central slice through a sub-tomogram average of PL sub-complexes in *A. tumefaciens*. Scale bar is 20 nm. **e, f and g**) Same as in (a, b and c) but for *H. neptunium*. Scale bars are 100 nm **h**) Central slice through a sub-tomogram average of PL sub-complexes in *H. neptunium*. Scale bar is 10 nm. **i, j and k**) Same as in (a, b and c) but for *C. crescentus*. Scale bars are 50 nm. **l**) Central slice through a sub-tomogram average of PL sub-complexes in *C. crescentus*. Scale bar is 10 nm. OM=outer membrane, IM= inner membrane.

**Figure 4:**
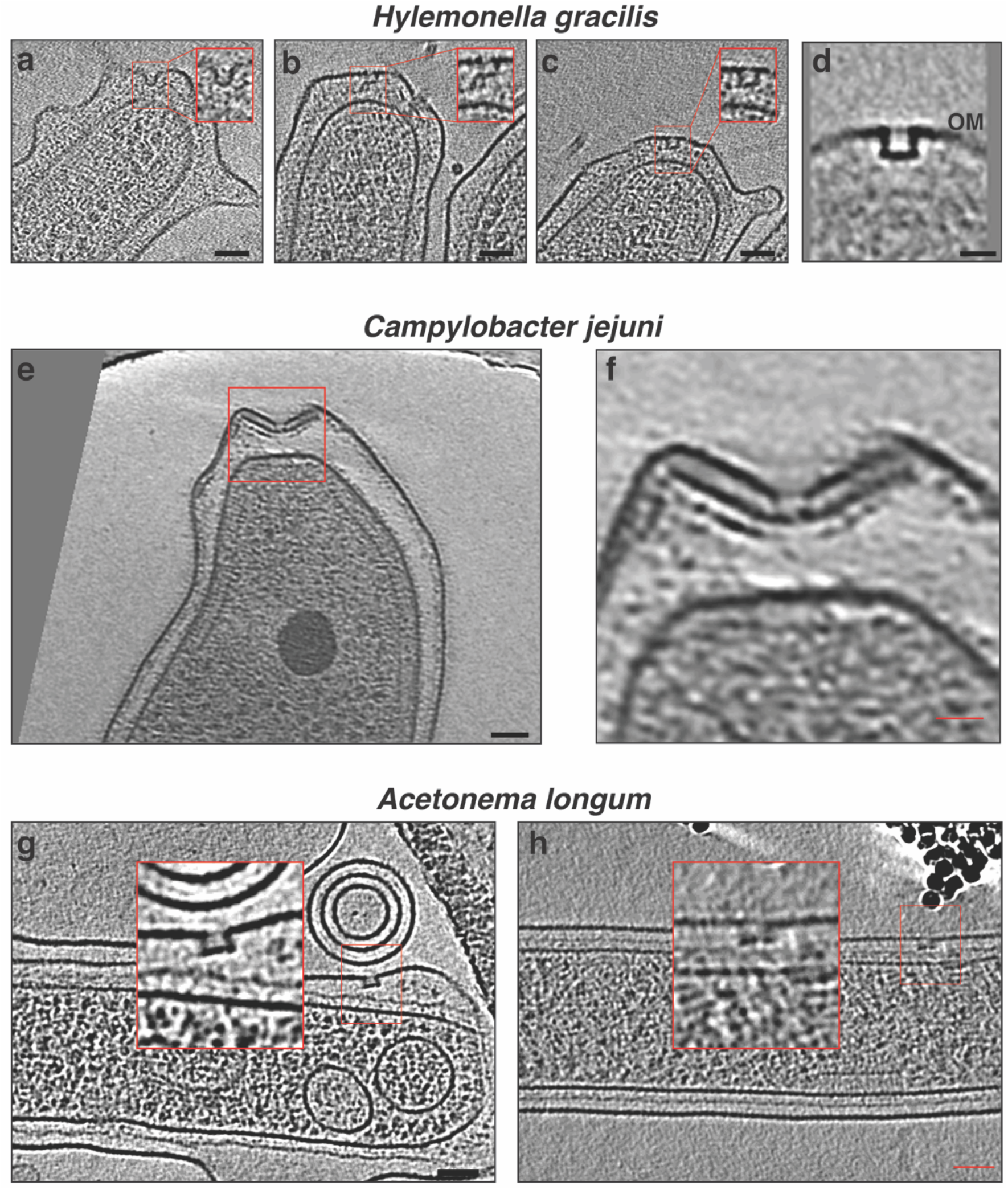
Cryo-ET of Betaproteobacteria, Epsilonproteobacteria and Firmicutes. **a, b and c**) Slices through electron cryo-tomograms of *H. gracilis* highlighting the presence of flagellar disassembly PL sub-complexes with zoom-ins of these sub-complexes present in the red squares. Scale bars are 50 nm. **d**) Central slice through a sub-tomogram average of PL sub-complexes in *H. gracilis*. Scale bar is 20 nm. **e**) A Slice through electron cryo-tomograms of *C. jejuni* highlighting the presence of a flagellar disassembly PL sub-complex (red square). Scale bars is 50 nm. **f**) A zoom-in of the area enclosed in the red square in e. Scale bar is 20 nm **g and h**) Slices through electron cryo-tomograms of *A. longum* highlighting the presence of flagellar disassembly PL subcomplexes with zoom-ins of these sub-complexes present in the red squares. Scale bars are 50 nm.

Having established that PL sub-complexes are widespread in Proteobacteria, we next looked for them in *Acetonema longum*, a diderm belonging to the class of Clostridia in the Firmicutes phylum. PL sub-complexes were found in *A. longum* as well (Fig. 4 g-h).

The presence of PL sub-complexes in diverse bacterial phyla could be because it is an ancient and conserved feature, or because the P- and L-ring proteins were recently horizontally transferred. To explore these possibilities, we performed an implicit phylogenetic analysis on all species in which PL sub-complexes have been found (by cryo-EM, 15 in total including the species described here plus those in Refs. (7, 16, 17)). We compared the sequence distances amongst FlgI’s (P-ring protein) and amongst FlgH’s (L-ring protein) as well as 25 single-copy well-conserved proteins (as previously described in Ref. (23)). This allowed us to investigate how P- and L-ring proteins evolved compared to the reference 25 proteins (24). If the sequence distances amongst FlgI (or FlgH) proteins in two species is smaller than the 25 reference proteins, this indicates a horizontal gene transfer event (24). This analysis of pairwise comparisons of the investigated species showed that the sequence distances between FlgH proteins is at least as divergent as the 25 reference proteins, and therefore there is no evidence of horizontal gene transfer between these species (Fig. 5 a and Table S1). This same result was seen for FlgI (Fig. 5 b and Table S2).

**Figure 5:**
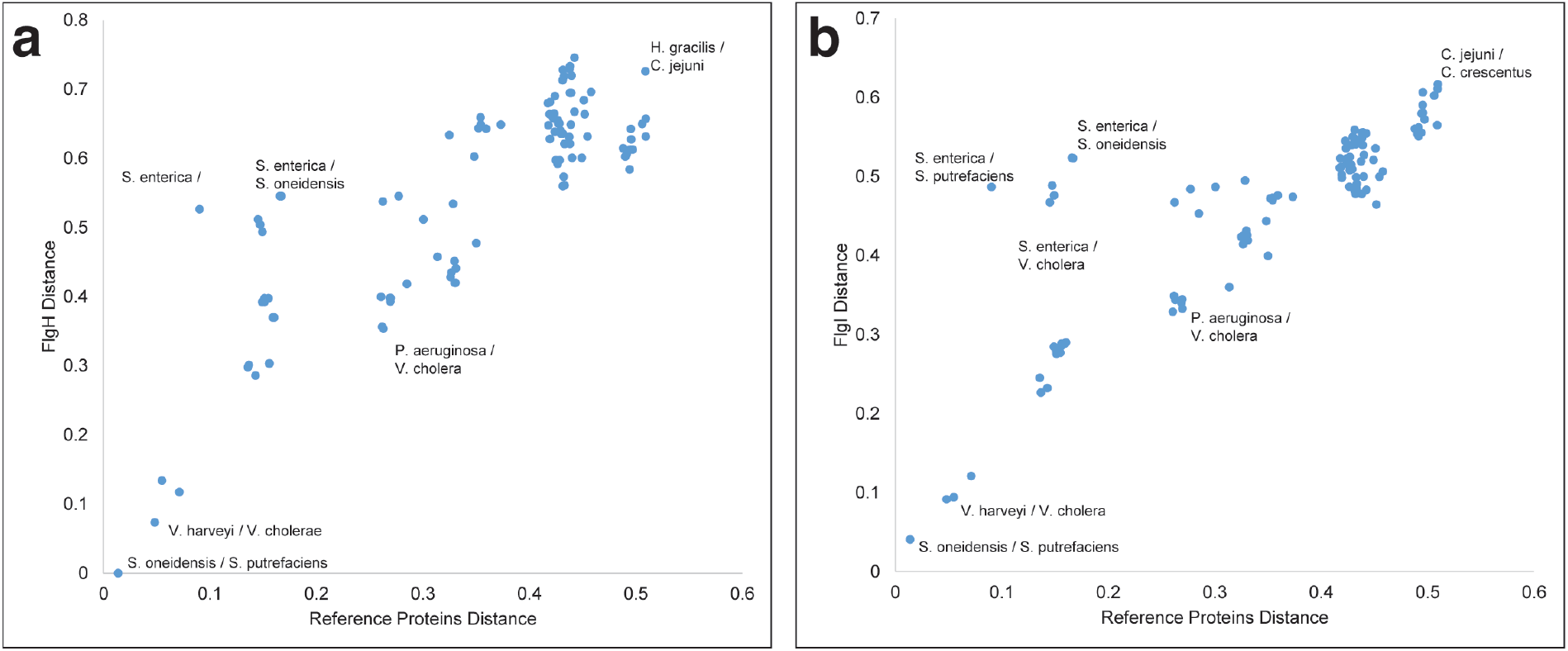
Implicit phylogenetic analysis of bacterial L- and P-rings protein. **a**) A scatter plot of pairwise sequence distance of the fifteen investigated species in this study based on concatenated 25 reference proteins and the L-ring protein, FlgH. Some examples of pairwise species comparisons are annotated in the plot for the sake of clarity. **b)** Same as in (**a**) but with the P-ring protein, FlgI. Plots shown in **a** and **b** are made with the primary copies of *P. shigelloides* and *S. putrefaciens* FlgI and FlgH proteins. For similar plots with the secondary copies of FlgI and FlgH in these two species see figures S4 and S5. The X- and Y-axes in these plots have arbitrary units.

In *Shewanella putrefaciens* and *Plesiomonas shigelloides* two copies for FlgI and FlgH were annotated. For both species and both genes, one copy showed more similarity to the nearest relative *(S. putrefaciens* FlgI: A4Y8M8, FlgH: A4Y8M9; *P. shigelloides* FlgI: R8AUG5, FlgH: R8AUH3, referred to as the primary copy). On the other hand, the other copy (referred to as secondary copy) showed more divergence to any studied organism *(S. putrefaciens* FlgI: A4YB38, FlgH: A4YB39; *P. shigelloides* FlgI: R8AS48, FlgH: R8AS34, see Figs. S4 & S5 and Tables S3 & S4). While two copies of these genes existed for these organisms, no evidence of horizontal gene transfer was present amongst the studied species implying that one of the copies could be due to a horizontal gene transfer from another species not included in this study or is a result of a gene duplication event.

## Discussion

An important step in reconstructing the evolutionary history of biomolecular complexes is to know when certain features and functions originated. Recent studies indicate that the bacterial flagellum is an ancient machine that originated from a single or few proteins through multiple gene duplication and diversification events that proceeds the common ancestor of bacteria (23). Some parts of the flagellar motor are homologous to other sub-complexes present in other machines. The stator proteins MotA/B are homologous to proteins in the Tol-pal and TonB systems while the motor’s ATPase is homologous to the beta subunit of the ATP synthase (23, 25). This suggests that other, even older machines donated features and functions to the first motor. Moreover, the Type III secretion system (T3SS), also known as the injectisome, is homologous to the bacterial flagellar motor (though the P- and L-rings of the motor are not homologous to the secretin part of the injectisome) (26). Because motility proceeded the evolution of eukaryotic cells, the targets of T3SS, and the T3SS is restricted mainly to proteobacteria, the T3SS likely derived from the flagellum (27, 28).

The proteins that form the P- and L-rings, namely FlgI and FlgH respectively, are present widely in flagellated bacteria, however, they are not as universal as other flagellar proteins known as the core proteins. For example, Spirochaetes (characterized by periplasmic flagella) and Firmicutes do not necessarily have the P- and L-rings. These two phyla are usually considered amongst the earliest evolved phyla of bacteria (29), indicating that although the P- and L-rings appeared early during the motor evolution, they were probably not present at first (23). The P- and L-rings have been thought to act as bushings supporting the rotation of the rod. The discovery that they persist after flagellar disassembly in an altered, sealed form, suggested an additional function – perhaps they remain to seal what would otherwise be a hole in the outer membrane. Here we have found that PL sub-complexes are widespread amongst Bacteria and ancient (not the result of recent horizontal gene transfers). This indicates that the putative outer-membrane-sealing function is important enough to have been conserved since the diversification of bacterial phyla.

In addition, we showed that in species with sheathed flagella, the outer membrane remained intact above PL sub-complexes, but the base of the PL sub-complexes was nevertheless apparently sealed. This raises questions about the nature and function of the PL sub-complex in these species. Does it serve a function distinct from membrane-sealing in *Vibrio*, or it could be a vestige retained in their evolution from ancestors with unsheathed flagella? Finally, it will be interesting to find out whether membrane seals are needed only for flagellar motor disassembly or if they might be needed in other closely related systems like the injectisome.

## Acknowledgements

This work was supported by NIH grant R35 GM122588 to GJJ. Cryo-EM work was done in the Beckman Institute Resource Center for Transmission Electron Microscopy at Caltech. M.K. acknowledges a Rubicon postdoctoral fellowship from De Nederlandse Organisatie voor Wetenschappelijk Onderzoek (NWO). Ariane Briegel kindly helped in collecting part of the data. We thank Catherine M. Oikonomou for reading the manuscript and for the insightful discussions. We thank Dr. Pat Zambryski from the University of California, Berkeley for providing us with *A. longum* strain used in this study.

## Supplementary Figures

**Figure S1:**
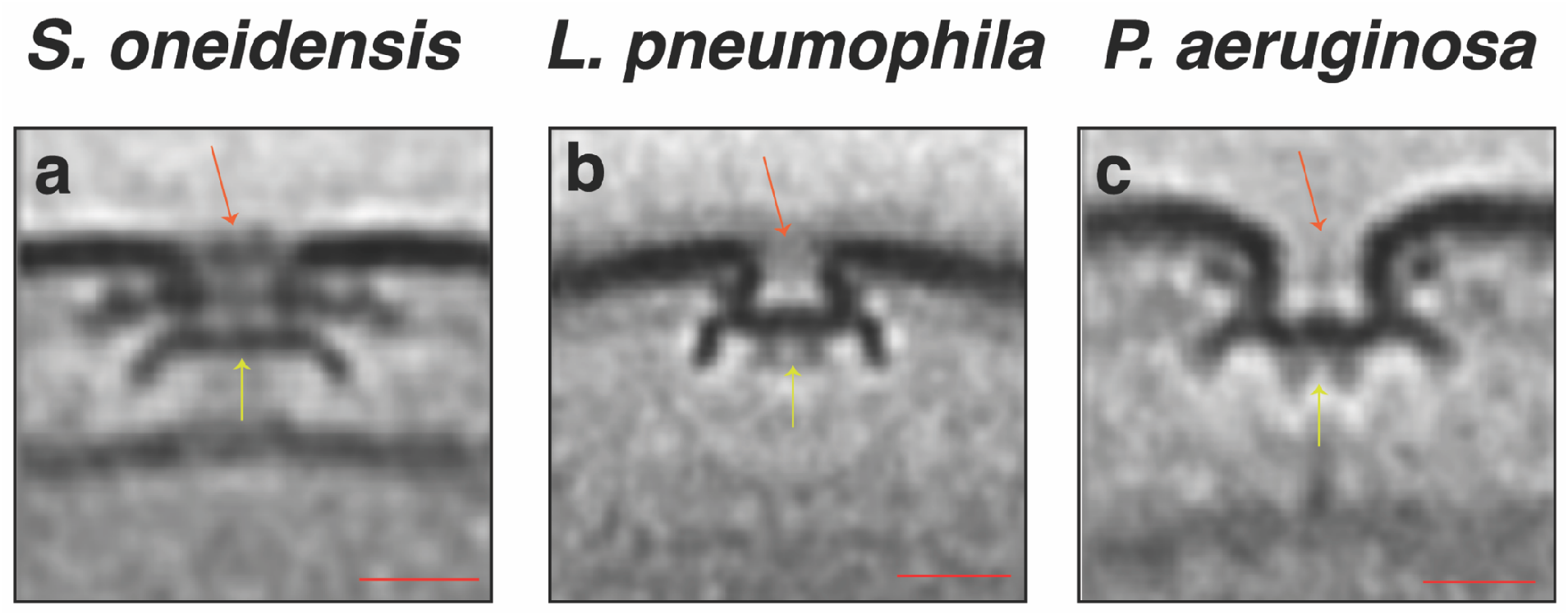
Central slices through sub-tomogram averages of PL sub-complexes in *S. oneidensis* (**a**), *L. pneumophila* (**b**) and *P. aeruginosa* (**c**). Scale bar is 20 nm. Orange arrows indicate the discontinuity in the outer membrane. Note the presence of two densities below the orange arrow in *S. oneidensis*. Yellow arrows point to the plug densities in these structures. Scale bars are 20 nm. These structures are adapted from reference (7).

**Figure S2:**
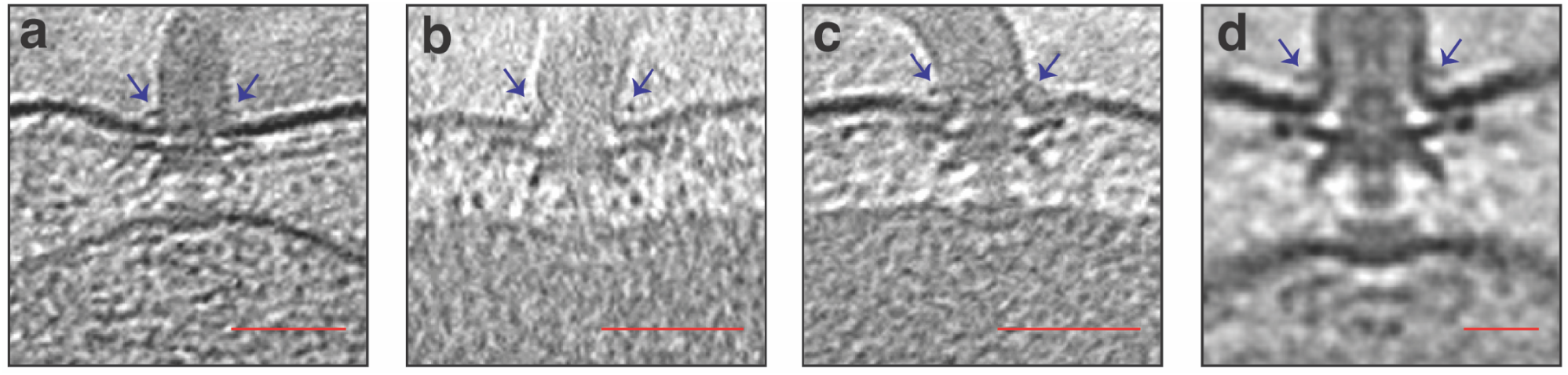
**a, b and c**) Slices through electron cryo-tomograms of *V. harvyei* with the blue arrows highlighting the presence of an extracellular ring at the bending of the outer membrane to form the sheath that surrounds the flagellar filament. Scale bars are 50 nm. **d**) A central slice through sub-tomogram average of the sheathed flagellar motor of *V. hanyei* obtained by averaging five particles only to indicate the presence of the extracellular ring (blue arrows). Scale bar is 20 nm.

**Figure S3:**
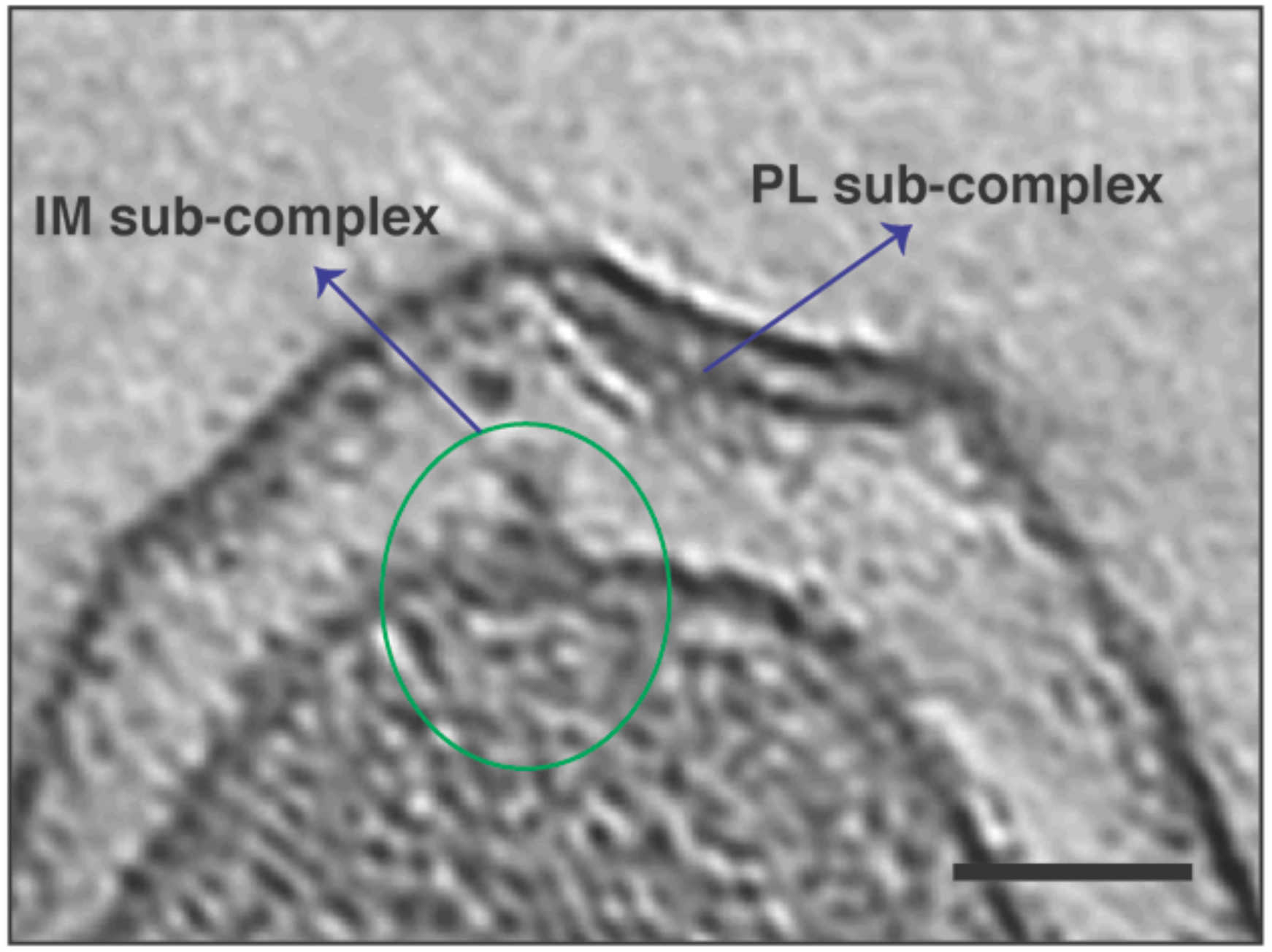
A slice through electron cryo-tomogram of a *C. jejuni* cell illustrating the presence of an inner-membrane (IM) associated sub-complex (green circle) next to the outer-membrane associated PL sub-complex. This is a diffrenet slice of the same example shown in Figure 4 e and f. Note that a similar observation has been recently described for *L. pneumophila* (7). Scale bar is 50 nm.

**Figure S4:**
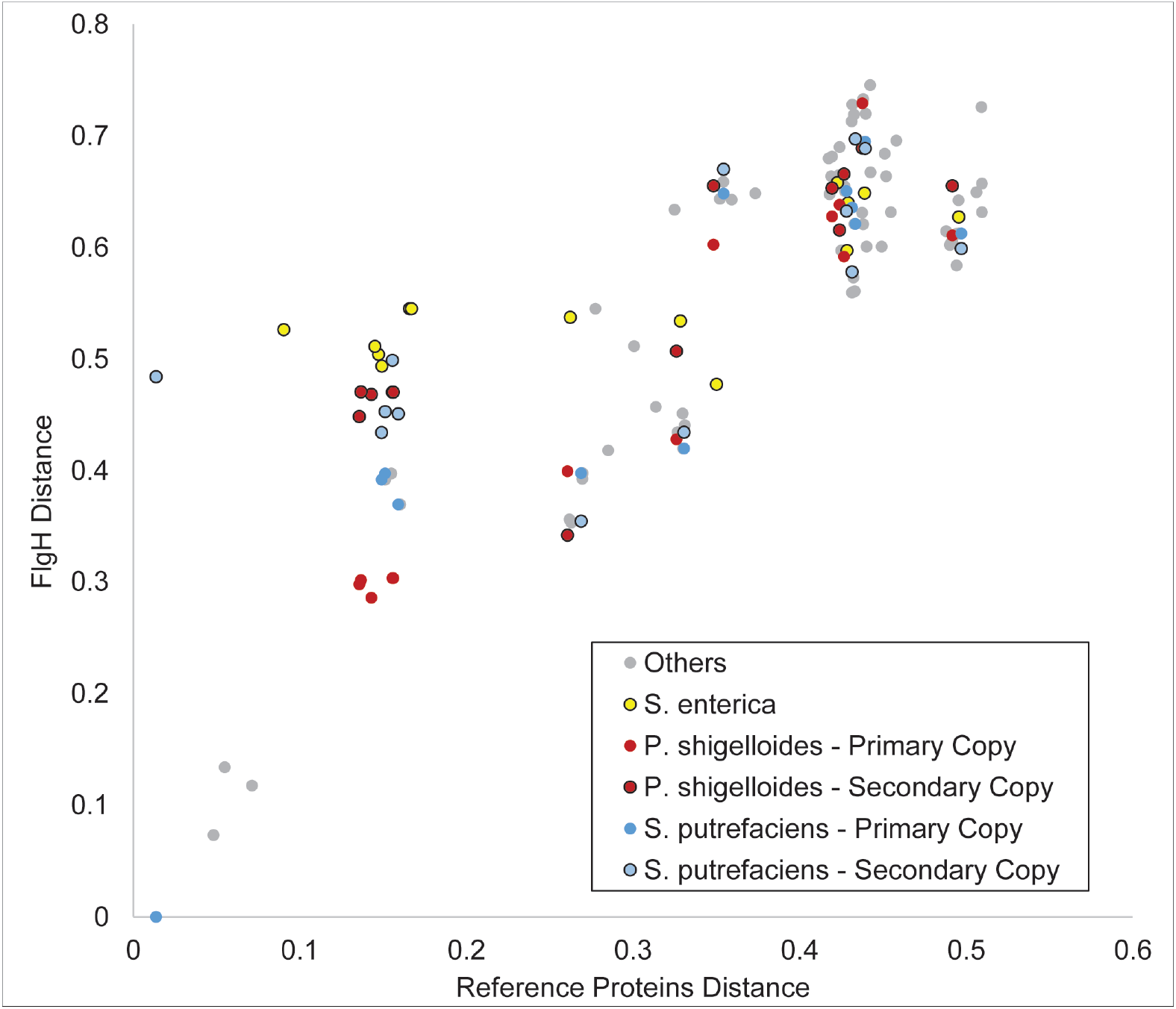
A scatter plot of the pairwise sequence distance of the investigated species based on concatenated 25 reference proteins and the L-ring protein, FlgH. In this plot both copies of FlgH proteins found in *S. putrefaciens* and P. *shigelloides* are used and highlighted. Interestingly, *Salmonella* FlgH protein is more divergent than expected based on the concatenated reference proteins distance. Note that in Figure 5a only the primary copies of *S. putrefaciens* and *P. shigelloides* are used. The X- and Y-axes in this plot have arbitrary units.

**Figure S5:**
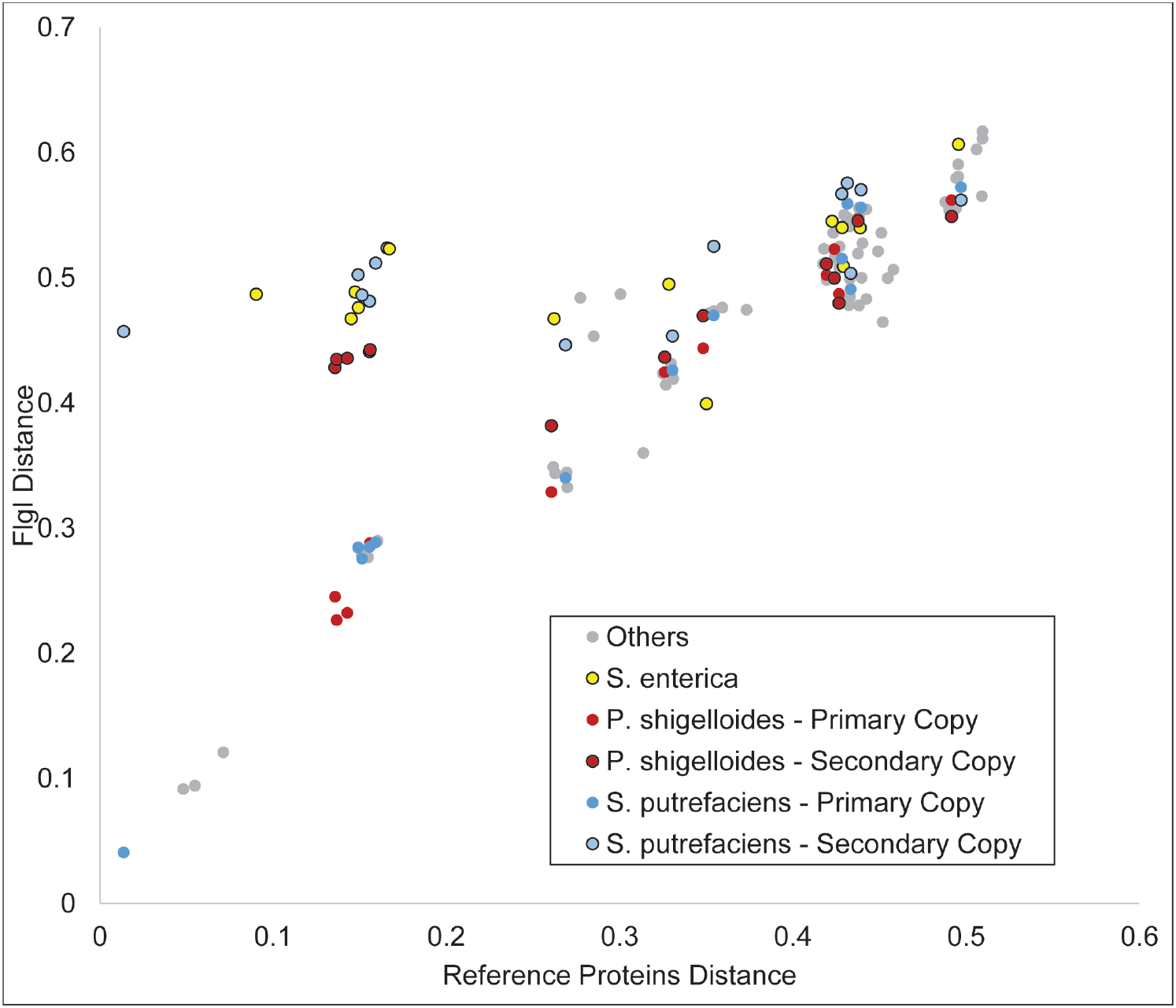
A scatter plot of the pairwise sequence distance of the investigated species based on concatenated 25 reference proteins and the P-ring protein, FlgI. In this plot both copies of FlgI proteins found in *S. putrefaciens* and *P. shigelloides* are used and highlighted. Interestingly, *Salmonella* FlgI protein is more divergent than expected based on the concatenated reference proteins distance. Note that in Figure 5b only the primary copies of *S. putrefaciens* and P. *shigelloides* are used. The X- and Y-axes in this plot have arbitrary units.

## Materials and Methods

### Cell types and growth conditions

*Vibrio cholerae* was grown 24 hours in LB at 30° C; diluted 150 μL into 2 mL Ca-HEPES buffer and grown at 30 °C for another 16 hours. *Vibrio harveyi* was grown in AB medium overnight at 30° C. *Vibrio fischeri* was grown overnight at 28° C in salt-supplemented LB medium with 35 mM MgSO4 (as described in (30)). Wild type *A. tumefaciens* C58 was transformed with Ti plasmid encoding for VirB8 fluorescently-tagged with GFP. Cells were grown overnight in LB at 28°C and subsequently spun down and resuspended to OD600=0.1 in AB medium supplemented with 300ug/ml streptomycin and100ug/ml spectinomycin. The cells were switched to 19°C and grown for 5h. To induce expression of VirB8-GFP, 200uM acetosyringone was added and cells were grown for 24h at 19°C. *H. neptunium* ATCC 15444 228405 cells were grown overnight in Marine Broth (MB) at 30° C. *C. crescentus* NA1000 565050 cells were synchronized in M2 buffer to get swarmer cells as described in references (31, 32). *C. jejuni* subsp. jejuni 81116 407148 were grown as described in reference (30). Briefly, cells were grown under microaerobic conditions for 48-60 hours on MH agar using CampyPak sachets (Oxoid) at 37° C. After that, cultures were restreaked and incubated for extra 16h. Then, bacteria were resuspended into 1 mL MH broth to an OD_600_ of 10 and were subsequently plunge-frozen. *H. gracilis* cells were grown for 48 hours in Broth 233 at 26°C without antibiotics to OD_600_ < 0.1,. Subsequently, cells were spun down at 1000 x *g* for 5 min and concentrated by ~10x for plunge freezing. *A. longum* were grown anaerobically on rhamnose as described in (33). Note that some of the tomograms were grown for other purposes other than observing their flagellar biogenesis.

### Cryo-ET sample preparation and imaging

10- or 20-nm gold beads were first coated with BSA and then the solution was mixed with cells. 3-4 μL of this mixture was applied to a glow-discharged, carbon-coated, R2/2, 200 mesh copper Quantifoil grid (Quantifoil Micro Tools) in a Vitrobot chamber (FEI) with 100% humidity at room temperature. Samples were blotted using Whatman paper and then plunge-frozen in ethane/propane mix. Imaging was done on an FEI Polara 300-keV field emission gun electron microscope (FEI company, Hillsboro, OR, USA) equipped with a Gatan image filter and K2 Summit direct electron detector in counting mode (Gatan, Pleasanton, CA, USA). Data were collected using the UCSF Tomography software (34) with each tilt series ranging from −60° to 60° in increments ranging from 1°-3°, and an underfocus range of ~5–10 μm for the different samples. A cumulative electron dose of 200 e^-^/A^2^ for each individual tilt series in *A. longum*, 200 e^-^/A^2^ for *A. tumefaciens*, 200 e^-^/A^2^ for *C. crescentus*, 75 e^-^/A^2^ for *H. gracilis*, 160 e^-^/A^2^ for *V. cholera*, 160 e^-^/A^2^ for *V. harveyi*, 150 e^-^/A^2^ for *V. fischeri*, 200 e^-^/A^2^ *H. neptunium*, 200 e^-^/A^2^ for *C. jejuni*.

### Image processing and subtomogram averaging

Three dimensional reconstructions of the tilt series were either done through automatic RAPTOR pipeline used in the Jensen lab at Caltech or by using the IMOD software package (35). Sub-tomogram averages with 2-fold symmetrization along the particle Y-axis were produced using PEET program (36). The number of PL sub-complexes that were averaged for each species are the following: 47 particles were averaged for the *V. cholera*, 4 particles for *V. harveyi*, 4 particles for *V. fischeri*, 6 particles for *A. tumefacienns*, 4 particles for *H. neptunium*, 5 particles for *C. crescentus*, 8 particles for *H. gracilis*.

### Bioinformatics analysis

An implicit phylogenetic approach was employed to detect the presence or absence of lateral gene transfer of flgI or flgH between sub-phylum of proteobacteria. In this analysis, species distance was estimated from the protein sequence distance between a set of single-copy cluster of orthologous genes (COGs) and gene distance was estimated from the distance between individual flagellar protein sequences. The set of single-copy COGs was taken from reference (29) and further refined to only 25 COGs that contained a single copy in all 15 species considered here. These COGs along with the flagellar proteins flgI and flgH were individually aligned with MUSCLE (37) with 100 maxiters. Conserved blocks were identified using Gblocks (38) with a maximum of 8 contiguous non-conserved positions, a minimum length of 2 for a block, half gap positions allowed, and a similarity matrix was employed. Following the individual processing of the single-copy COGs, the individual multiple sequences alignments (MSA) were concatenated to create a species-level alignment. Pairwise distances within the MSA of flagellar protein sequences and within the MSA of concatenated single-copy COGs were calculated using the DistanceMatrix library in Biopython with the BLOSUM62 substitution matrix.

### Constructing the taxonomic tree

400 Representative bacterial species were selected at random from all bacteria in UniProt with a reference proteome annotation. Species included in this study were appended to this list. The taxonomic tree was rendered using ETE (39).

## References

1. R. M. Macnab, How Bacteria Assemble Flagella. Annual Review of Microbiology 57, 77–100 (2003).

2. C. J. Jones, R. M. Macnab, Flagellar assembly in Salmonella typhimurium: analysis with temperature-sensitive mutants. Journal of Bacteriology 172, 1327–1339 (1990).

3. T. Kubori, N. Shimamoto, S. Yamaguchi, K. Namba, S.-I. Aizawa, Morphological pathway of flagellar assembly in Salmonella typhimurium. Journal of Molecular Biology 226, 433–446 (1992).

4. H. Li, V. Sourjik, Assembly and stability of flagellar motor in Escherichia coli: Flagellar motor assembly. Molecular Microbiology 80, 886–899 (2011).

5. F. D. Fabiani, et al., A flagellum-specific chaperone facilitates assembly of the core type III export apparatus of the bacterial flagellum. PLOS Biology 15, e2002267 (2017).

6. E. J. Cohen, K. T. Hughes, Rod-to-hook transition for extracellular flagellum assembly is catalyzed by the L-ring-dependent rod scaffold removal. J. Bacteriol. 196, 2387–2395 (2014).

7. M. Kaplan, et al., *In situ* imaging of the bacterial flagellar motor disassembly and assembly processes. The EMBO Journal, e100957 (2019).

8. S. Chen, et al., Structural diversity of bacterial flagellar motors: Structural diversity of bacterial flagellar motors. The EMBO Journal 30, 2972–2981 (2011).

9. M. Kaplan, et al., The presence and absence of periplasmic rings in bacterial flagellar motors correlates with stator type. eLife 8 (2019).

10. Z. Qin, W. Lin, S. Zhu, A. T. Franco, J. Liu, Imaging the Motility and Chemotaxis Machineries in Helicobacter pylori by Cryo-Electron Tomography. Journal of Bacteriology 199, e00695–16 (2017).

11. X. Zhao, S. J. Norris, J. Liu, Molecular Architecture of the Bacterial Flagellar Motor in Cells. Biochemistry 53, 4323–4333 (2014).

12. B. Chaban, I. Coleman, M. Beeby, Evolution of higher torque in Campylobacter-type bacterial flagellar motors. Scientific Reports 8 (2018).

13. H. Terashima, H. Fukuoka, T. Yakushi, S. Kojima, M. Homma, The Vibrio motor proteins, MotX and MotY, are associated with the basal body of Na^+^-driven flagella and required for stator formation. Molecular Microbiology 62, 1170–1180 (2006).

14. H. Terashima, M. Koike, S. Kojima, M. Homma, The Flagellar Basal Body-Associated Protein FlgT Is Essential for a Novel Ring Structure in the Sodium-Driven Vibrio Motor. Journal of Bacteriology 192, 5609–5615 (2010).

15. J. M. Skerker, M. T. Laub, Cell-cycle progression and the generation of asymmetry in Caulobacter crescentus. Nature Reviews Microbiology 2, 325–337 (2004).

16. J. L. Ferreira, et al., γ-proteobacteria eject their polar flagella under nutrient depletion, retaining flagellar motor relic structures. PLOS Biology 17, e3000165 (2019).

17. S. Zhu, et al., *In Situ* Structures of Polar and Lateral Flagella Revealed by Cryo-Electron Tomography. Journal of Bacteriology 201 (2019).

18. X.-Y. Zhuang, et al., Dynamic production and loss of flagellar filaments during the bacterial life cycle. bioRxiv (2019) https:/doi.org/10.1101/767319 (September 27, 2019).

19. C. M. Oikonomou, G. J. Jensen, A new view into prokaryotic cell biology from electron cryotomography. Nature Reviews Microbiology 15, 128 (2017).

20. J. A. G. Briggs, Structural biology in situ--the potential of subtomogram averaging. Curr. Opin. Struct. Biol. 23, 261–267 (2013).

21. K. E. Leigh, et al., “Subtomogram averaging from cryo-electron tomograms” in Methods in Cell Biology, (Elsevier, 2019), pp. 217–259.

22. S. Zhu, et al., Molecular architecture of the sheathed polar flagellum in *Vibrio alginolyticus*. Proceedings of the National Academy of Sciences, 201712489 (2017).

23. R. Liu, H. Ochman, Stepwise formation of the bacterial flagellar system. Proceedings of the National Academy of Sciences 104, 7116–7121 (2007).

24. M. Ravenhall, N. Škunca, F. Lassalle, C. Dessimoz, Inferring Horizontal Gene Transfer. PLOS Computational Biology 11, e1004095 (2015).

25. E. Cascales, R. Lloubès, J. N. Sturgis, The TolQ-TolR proteins energize TolA and share homologies with the flagellar motor proteins MotA-MotB: TolQ-TolR are needed to energize TolA. Molecular Microbiology 42, 795–807 (2008).

26. A. Diepold, J. P. Armitage, Type III secretion systems: the bacterial flagellum and the injectisome. Philosophical Transactions of the Royal Society B: Biological Sciences 370, 20150020 (2015).

27. M. Saier, Evolution of bacterial type III protein secretion systems. Trends in Microbiology 12, 113–115 (2004).

28. W. Deng, et al., Assembly, structure, function and regulation of type III secretion systems. Nature Reviews Microbiology 15, 323–337 (2017).

29. F. D. Ciccarelli, Toward Automatic Reconstruction of a Highly Resolved Tree of Life. Science 311, 1283–1287 (2006).

30. M. Beeby, et al., Diverse high-torque bacterial flagellar motors assemble wider stator rings using a conserved protein scaffold. Proceedings of the National Academy of Sciences 113, E1917–E1926 (2016).

31. J.-W. Tsai, M. R. K. Alley, Proteolysis of the Caulobacter McpA Chemoreceptor Is Cell Cycle Regulated by a ClpX-Dependent Pathway. Journal of Bacteriology 183, 5001–5007 (2001).

32. A. Briegel, et al., Multiple large filament bundles observed in Caulobacter crescentus by electron cryotomography. Molecular Microbiology 62, 5–14 (2006).

33. E. I. Tocheva, et al., Polyphosphate storage during sporulation in the gram-negative bacterium Acetonema longum. J. Bacteriol. 195, 3940–3946 (2013).

34. S. Q. Zheng, et al., UCSF tomography: an integrated software suite for real-time electron microscopic tomographic data collection, alignment, and reconstruction. J. Struct. Biol. 157, 138–147 (2007).

35. J. R. Kremer, D. N. Mastronarde, J. R. McIntosh, Computer visualization of three-dimensional image data using IMOD. J. Struct. Biol. 116, 71–76 (1996).

36. D. Nicastro, The Molecular Architecture of Axonemes Revealed by Cryoelectron Tomography. Science 313, 944–948 (2006).

37. R. C. Edgar, MUSCLE: multiple sequence alignment with high accuracy and high throughput. Nucleic Acids Res. 32, 1792–1797 (2004).

38. G. Talavera, J. Castresana, Improvement of Phylogenies after Removing Divergent and Ambiguously Aligned Blocks from Protein Sequence Alignments. Systematic Biology 56, 564–577 (2007).

39. J. Huerta-Cepas, F. Serra, P. Bork, ETE 3: Reconstruction, Analysis, and Visualization of Phylogenomic Data. Molecular Biology and Evolution 33, 1635–1638 (2016).

